# Virulence, Aggressiveness, and Fungicide Sensitivity of *Phytophthora spp*. Associated with Soybean in Illinois

**DOI:** 10.1101/2022.06.30.498288

**Authors:** Daniel G. Cerritos-Garcia, Shun-Yuan Huang, Nathan M. Kleczewski, Santiago X. Mideros

## Abstract

Phytophthora root and stem rot (PRR), caused by *Phytophthora sojae*, is one of the most devastating oomycete diseases of soybean in Illinois. Single resistant genes (*Rps*) are used to manage this pathogen, but *P. sojae* has adapted to *Rps* causing failure of resistance in many regions. In addition to *P. sojae*, recent reports indicate that *Phytophthora sansomeana* could also cause root rot in soybean. Soil samples and symptomatic plants were collected across 40 Illinois counties between 2016 and 2018. *Phytophthora sojae* (77%) was more abundant than *P. sansomeana* (23%) across Illinois fields. Both species were characterized by virulence, aggressiveness, and fungicide sensitivity. Virulence of all isolates was evaluated using the hypocotyl inoculation technique in 13 soybean differentials. Aggressiveness was evaluated in the greenhouse by inoculating a susceptible cultivar and measuring root and shoot dry weight. On average, *P. sojae* isolates were able to cause disease on six soybean differentials. *P. sojae* was more aggressive than *P. sansomeana*. All isolates were sensitive to azoxystrobin, ethaboxam, mefenoxam, and metalaxyl. The characterization of the population of species associated with PRR will inform management decisions for this disease in Illinois.

## Introduction

Soybean production in Illinois and the U.S. is severely affected by Phytophthora root and stem rot (PRR) caused by the oomycete *Phytophthora sojae* Kaufmann & Gerdemann (Bradley et al. 2021). PRR has been the fourth most destructive soybean disease in the state for two decades (1996 to 2016), causing estimated losses of 945 million U.S. dollars during this period (Bandara et al. 2020). *P. sojae* is a soil-borne pathogen that can infect soybean at any growth stage, causing damping-off in seedlings and root and stem rot in older plants (Schmitthenner 2000). The use of cultivars with single resistance genes to *P. sojae* (*Rps*) has been the primary method of management, but the pathogen has adapted to various deployed genes (*Rps*1a, *Rps*1c, *Rps*1k, *Rps*3a, and *Rps*6) thus rendering them ineffective in some regions of the Midwest U.S. (Dorrance et al. 2018).

To monitor changes in virulence towards *Rps* genes and *P. sojae* pathotype diversity, surveys have been conducted across different states in the Midwest U.S. (Dorrance et al. 2016; Dorrance et al. 2018). In these surveys, pathotypes of *P. sojae* were identified based on their virulence to differentials carrying single *Rps* genes (Dorrance et al. 2008). Throughout the years, pathotypes and the proportion of isolates with virulence to the most commonly deployed resistance genes, *Rps*1k and *Rps*1c, have increased (Malvick and Grunden 2004; Dorrance et al. 2016). In addition, the virulence of *P. sojae* to *Rps* genes measured as isolate complexity (the number of *Rps* genes that an isolate is virulent to) also continue to increase (Dorrance et al. 2016).

The increase in pathotype complexity makes long-term management of this disease challenging. However, there are evolutionary constraints that might eventually limit the fitness of virulent strains (Lannou 2012). Of particular importance for plant disease management are the costs of virulence. One method to identify costs of virulence is to measure the aggressiveness (defined as the quantitative variation in pathogenicity on susceptible hosts) of pathogen strains (Zhan and McDonald 2013). Fitness costs, such as lower spore production, due to increased virulence have been previously reported for many pathogens including the oomycete *P. infestans* (Montarry et al. 2010), and the soil inhabitant *Leptosphaeria maculans* (Daverdin et al. 2012). Fitness related traits such as the aggressiveness of *P. sojae* isolates have not been previously reported. A better knowledge of the relationships between virulence and aggressiveness in *P. sojae* populations can be used to make suggestions for deployment of major resistance genes (Zhan and McDonald 2013).

In addition to *P. sojae*, a second species, *Phytophthora sansomeana* E.M. Hansen & Reeser, has been reported as pathogenic to soybean (Hansen et al. 2009; Rojas et al. 2017a). Unlike *P. sojae, P. sansomeana* has a broad host range and can infect corn, alfalfa, peas, apples, pears, Douglas fir, and Fraser fir (Hansen et al. 2009; Zelaya-Molina et al. 2009; Chang et al. 2017; Farr and Rossman 2022). Infection by *P. sansomeana* causes damping-off and root rot, but not stem rot, which is typical of *P. sojae* (Hansen et al. 2009; Rojas et al. 2017b). Although *P. sansomeana* was formally described in 2009, a second *Phytophthora* sp. causing PRR was first reported in Illinois in 2004 (Malvick and Gruden 2004). In a survey of oomycete pathogens causing soybean seedling diseases, Rojas et al. (2017a) isolated *P. sansomeana* from Illinois and five other U.S. states and found it was among the most aggressive species. Little is known about the prevalence of *P. sansomeana* in Illinois soils, its contribution to PRR, and if the current management practices for *P. sojae* would effectively manage *P. sansomeana*.

In addition to *Rps* genes, PRR can be managed with quantitative host resistance and fungicide seed treatments (Dorrance et al. 2009; Dorrance 2018; Cerritos-Garcia et al. 2021). Quantitative resistance (also known as partial resistance) is not race-specific, and therefore broadly effective against all pathotypes of *P. sojae* (Dorrance et al. 2003). However, quantitative resistance is expressed after emergence after the production of the first unifoliate leaves, hence increasing the likelihood for seedling blight, especially in fields where a large portion of the population of *P. sojae* are insensitive to the *Rps* gene contained within the crop. This is one reason combining resistance with a PRR labeled fungicide seed treatment is recommended for effective management (Dorrance and McClure 2001).

For decades, the phenylamides [Fungicide Resistance Action Committee (FRAC) code 4], mefenoxam, and metalaxyl, have been the primary fungicide active ingredients used in seed reatments to control *P. sojae* (Dorrance 2018). Some Quinone outside Inhibitors (QoI; FRAC code 11) fungicide active ingredients, for example, azoxystrobin, have limited efficacy on *P. sojae* (Radmer et al 2017; Crop Protection Network 2020). More recently, the active ingredients ethaboxam (FRAC 22) and oxathiapiprolin (FRAC 49) have been registered for use in soybean for oomycetes (Radmer et al. 2017). Little is known about the performance of these fungicide active ingredients on the Illinois populations of *P. sojae* or other Phytophthora that may be associated with soybean in the state.

Surveys assessing pathotype diversity among *P. sojae* populations are critical in evaluating the effectiveness of resistance genes. A population genetic study for a subset of the isolates reported here along with isolates from surrounding states presented evidence for population differentiation, with the most distinct isolates found in Illinois (Hebb et al. 2022). The objectives of this study were to characterize Phytophthora species associated with soybeans in Illinois by assessing isolate: i) virulence; ii) aggressiveness, and iii) sensitivity to PRR-labeled fungicide active ingredients. We evaluated the relationship between isolate aggressiveness and virulence. Characterizing pathogens that contribute to soybean’s root and stem rot will inform management decisions for this disease within the state.

## Materials and methods

### Sample collection

In fall 2016 and spring 2017, 50 soil samples were collected from 26 counties (Figure 1). Some fields were previously sampled (Malvick and Grunden, 2004; Dorrance et al. 2016) while others were chosen because of accessibility. In each field, an area of about 7.62 × 53.4 m with favorable conditions for oomycetes (poorly drained, high humidity) was selected, and ten subsamples were collected following a zigzag pattern. Subsamples were collected at a depth of 15 cm and placed in a single collection bag per field. Soil samples were dried at room temperature and then stored in a cold room at 4°C until processing.

**Figure 1.**
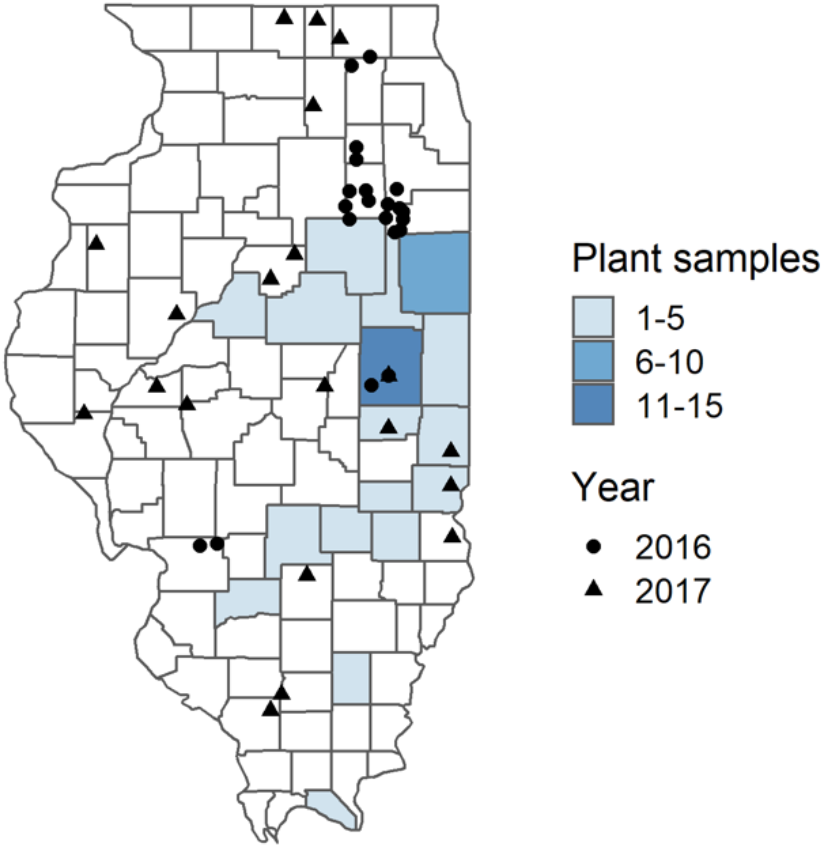
Location of soil samples in 2016 (circles) and 2017 (triangles), and counties from where symptomatic plants were received in 2018 (filled counties).

In the summer of 2018, diseased soybean plants from 51 fields in 19 counties were received from the Plant Diagnostic Clinic at the University of Illinois at Urbana-Champaign (Figure 1). All plants tested positive for the Phytophthora ImmunoStrip assay (Agdia, Elkhart, IN). Samples were stored in a cold room at 4°C and processed within two weeks.

### Soil baiting

Soil samples were ground using a Dynacrush soil mill (DC-5, Custom Laboratory Equipment Inc. Orange City, FL) and then subdivided into three to 10 plastic pots (15 cm diameter) (Dorrance et al. 2008). The soil in plastic pots was saturated with deionized, non-chlorinated water in the greenhouse, left overnight at 24-27°C, and drained the next day. After 24 to 48 hours, pots were placed in plastic bags and incubated in the dark at room temperature (∼22°C). After incubation, pots were moved to the greenhouse, and 15 to 20 surface-sterilized soybean seeds (immersion in 0.05% NaClO for 30 sec and then rinsed with sterile water) of the susceptible cultivar Sloan (no *Rps* gene and low partial resistance) were planted in the soil. Three days after planting, the pots were again saturated with water when seeds germinated. Two weeks later, symptomatic seedlings that presented brown to tan lesions were selected for isolation.

### Isolates

For isolation of *Phytophthora* spp., symptomatic seedlings were washed with sterilized, distilled water (Dorrance et al. 2008). Then, symptomatic tissue was cut into 1 to 2-cm sections, surface-sterilized for 10 seconds in 0.5% NaClO, and washed with sterilized water. The symptomatic tissue was dried on a sterilized paper towel and plated on PBNIC selective media (Dorrance et al. 2008). Petri plates were incubated at room temperature in the dark and monitored daily for mycelial growth. Isolates with coenocytic mycelia were transferred to either lima bean agar (LBA) or V8 juice agar. On these plates a preliminary morphological identification was conducted based in the size, shape and form of the oogonia. Isolates classified as oomycetes (based on morphology) were stored in vials with V8 juice agar at 15°C for molecular identification. Oomycetes were isolated from symptomatic plants received from the Plant Clinic in 2018 using the same procedure, except that plants were first washed with tap water and dish soap to remove soil particles.

### Isolate identification

An agar plug from each culture in the storage vials was transferred to an Erlenmeyer flask containing 125 to 150 ml of V8 broth, and isolates were cultured for one week on an orbital shaker (New Brunswick Scientific, Edison, NJ). After a week, approximately 100 mg of mycelia were transferred to FastDNA Lysing Matrix A tubes, and DNA was extracted using the FastDNA SPIN Kit protocol (MP Biomedicals, Solon, OH). Amplification of the ITS region utilized the ITS 4 and ITS 5 primer pair (White et al. 1990). The PCR product was cleaned using the Wizard SV Gel and PCR Clean-Up System (Promega, Madison, WI), and samples were submitted to the Roy J. Carver Biotechnology Center for Sanger sequencing using the ITS4 primer. The sequences were manually inspected, trimmed, and subjected to a BLAST alignment on the Phytophthora-ID website (http://phytophthora-id.org/) (Grünwald et al. 2011). All isolates recovered from soil samples were sequenced. The identity of the isolates collected from plant samples was assigned based on measurement of growth rate on PDA (Kaufmann and Gerdemann 1958), followed by morphological confirmation (Hansen et al. 2009).

### Virulence characterization

Pathotypes for each isolate were determined by inoculating 13 soybean differentials using the hypocotyl inoculation technique (Dorrance et al. 2008). The following differentials were used: Williams (universal susceptible, *rps*), Harlon (*Rps*1a), Harosoy13XX (*Rps*1b), Williams79 (*Rps*1c), Williams82 (*Rps*1k), L76-1988 (*Rps*2), L83-570 (*Rps*3a), PRX-146-36 (*Rps*3b), PRX-145-48 (*Rps*3c), L85-2352 (*Rps*4), L85-3059 (*Rps*5), Harosoy62XX (*Rps*6), and Harosoy (*Rps*7) (Dorrance et al. 2004). Twenty seeds of each differential were surface sterilized with 1% bleach for 30 seconds and planted in 1020-tray inserts filled with coarse vermiculite. After planting, to avoid fungal contamination, the trays were sprayed with a 0.6% benomyl solution (0.6 g Benlate 50 WP) and placed in the greenhouse at 25-28°C for a 16-hour photoperiod. Trays were watered and sprayed daily with the benomyl solution. One week after planting, 10 to 15 seedlings of each differential were selected for inoculation.

The inoculum was prepared as described by Dorrance et al. (2008). Briefly, *Phytophthora* spp. isolates were sub-cultured in LBA or V8 media and incubated at room temperature (∼22°C). After 7 days, each agar culture was transferred to a 10 ml syringe and macerated by forcing it through the syringe into another syringe with an 18-gauge needle used for inoculation. One-week-old soybean seedlings prepared as indicated above were inoculated by making a 1 cm slit in the hypocotyl with the 18-gauge needle and injecting 0.2 to 0.4 ml of culture slurry on and around the slit. The slit was then covered with Parafilm to maintain humidity. Inoculated seedlings were incubated in a humidity-controlled moisture chamber (95% humidity) for 48 hours at 20-22 °C in the dark, then transferred to the greenhouse (25-28°C, 16-hour photoperiod) evaluated for disease resistance after seven days. Reactions were scored as resistant when ≤ 25% of the seedlings were dead or susceptible when ≥ 75% were killed (Dorrance et al. 2008).

### Aggressiveness characterization

We used a modified version of the layer test to evaluate the aggressiveness of *Phytophthora* spp. isolates (Stewart and Robertson 2011; Rojas et al. 2017a). Eleven-centimeter-wide Jiffypots (Jiffy, Shippagan, Canada) with three holes at the bottom for drainage were used for the experiment. The pots were filled with 100 ml of coarse vermiculite, followed by the agar inoculum layer, and the second layer of 200 ml of vermiculite. Eight seeds of the susceptible cultivar, Sloan, were placed 2.5 cm over the inoculum and then covered with an additional 200 ml of vermiculite. Inoculum consisted of a double layer of a two-week-old *Phytophthora* spp. culture grown on V8 agar in Petri dishes (100 × 15 mm). We poured approximately 20 ml of V8 medium per Petri dish. The experiment was arranged in a completely randomized design with three replicates per treatment (isolates and controls) and pot as the experimental unit. Treatments included pots with a V8 agar layer with no *Phytophthora* spp. (mock control) and pots filled only with vermiculite (not-inoculated control). The experiment was repeated three times.

After planting, the pots were saturated with water until runoff and placed in the greenhouse (settings were 24°C and 16-hour photoperiod) and watered to capacity daily. Seedlings were removed from the pots, and the roots were washed with tap water to remove the vermiculite 14 days after germination. Shoots and roots were separated, placed in paper bags, and dried at 50°C in a laboratory oven (Precision Thelco oven, Thermo Fisher Scientific). The dry weight of shoots and roots was measured once consistent drymass had been reached (after 48 h of drying). The seedlings root and shoot dry weights were determined.

### Fungicide sensitivity

One *P. sojae* isolate from each county (*n* = 9) and all *P. sansomeana* isolates (*n* = 7) were assessed for fungicide sensitivity using poison plate assays. Isolates were screened against azoxystrobin (Syngenta Crop Protection, Greensboro, NC), ethaboxam (Valent U.S.A. LLC, Walnut Creek, CA), mefenoxam (Syngenta Crop Protection, Greensboro, NC), and metalaxyl (Gustafson, Plano, TX). Technical grade fungicides were dissolved in dimethyl sulfoxide (DMSO) to prepare stock solutions of 10 mg of a.i./ml. Then, we diluted stock solutions and added them to LBA medium (50-55°C) to obtain final concentrations of 0, 0.1, 0.5, 0.1, 1 and 10 μg/ml for azoxystrobin; 0, 0.001, 0.01, 0.05, 0.1 and 1 μg/ml for ethaboxam; and 0, 0.01, 0.05, 0.1, 0.5 and 1 μg/ml for mefenoxam and metalaxyl. One ml of DMSO was added to the medium as control (dose 0). For azoxystrobin, 25 mg of salicylhydroxamic acid (SHAM; Fisher Scientific) was added to the medium to inhibit the alternative oxidase pathway (Broders et al. 2007). The amended medium was poured into Petri dishes (60 × 15 mm). The concentrations for fungicide dilutions and SHAM were determined based on preliminary experiments.

*Phytophthora sojae* (*n* = 9) and *P. sansomeana* (*n* = 7) were grown in a V8 juice medium at room temperature as indicated above. After two weeks of growth, agar plugs (4mm diameter) of each isolate were transferred to the center of the fungicide amended plates. Plates were incubated at 25°C with a 10 h photoperiod (Precision Incubator Model 815, Thelco). Colony growth was determined after seven days by measuring colony diameter twice at right angles. Percent growth relative to the control was calculated by dividing the length of the colony of the fungicide amended plate by the average length of the control plates. Each isolate was replicated three times per concentration, and the experiment was repeated twice.

### Data analysis

The pathotype of each isolate, pathotype frequency, and complexity (the number of *Rps* genes that an isolate can overcome and cause disease) was determined using the package ‘hagis’ (McCoy et al. 2019) as implemented in R version 4.0.2. A linear mixed model was fit to the aggressiveness data (root dry weight and shoot dry weight) with species and isolates as fixed effects and experiment as a random effect. Models were fit using the package ‘lme4,’ and then the ANOVAs were conducted with the package ‘lmerTest’ (Bates et al. 2015; Kuznetsova et al. 2017). Residual plots were generated with the package ‘ggResidpanel’ to check model assumptions (Goode and Rey 2019). Estimated marginal means were obtained, and mean separation was done using Tukey’s HSD with the package ‘emmeans’ (Lenth 2020). For fungicide sensitivity data, the effective fungicide concentration to reduce the growth by 50% (EC50) was calculated for each isolate using the log-logistic four-parameter model in the package ‘drc’ (Ritz et al. 2015). The absolute EC_50_ was calculated using the “ED” function (Noel et al. 2018). Pearson correlation was performed to determine any relationship between pathotype complexity and aggressiveness. All the data and code used for analysis are available at https://github.com/danielcerritos/phytophthora.

## Results

### Identification of oomycetes from symptomatic soybean plants and soybean field soils in IL

A total of 148 isolates were recovered from soils and symptomatic soybean plants from 32 counties. The recovered isolates were categorized into 12 species based on morphology and ITS sequences. Overall, *Pythium* spp. dominated both field soils and diseased plants. *Pythium ultimum* var. *ultimum* (42%) was the most abundant species, followed by *P. sojae* (24%) and *P. sansomeana* (7%). The remaining nine species included: *Pythium yorkensis, Pythium pleroticum, Pythium aphanidermatum, Pythium torulosum, Pythium vexans, Pythium ultimum* var. *sporangiiferum, Pythium acanthophoron, Pythium irregulare*, and *Pythium acrogynum*. Characterization of the *Pythium* spp. population will be published elsewhere. A total of 31 isolates were identified as *Phytophthora* spp. (Table 1), which were recovered from eight of the 50 soil samples (16%) and eight of the 51 plant samples (16%). Phytophthora isolates were recoverd from a total of 40 samples collected from 12 counties. *Phytophthora sojae* and *P. sansomeana* were recovered from both soil and symptomatic plants. *Phytophthora sojae* was recovered from nine fields in 10 counties, with more than one isolate recovered from seven fields. *Phytophthora sansomeana* was recovered from five fields in five counties, and more than one isolate was recovered from two of these fields.

**Table 1.**
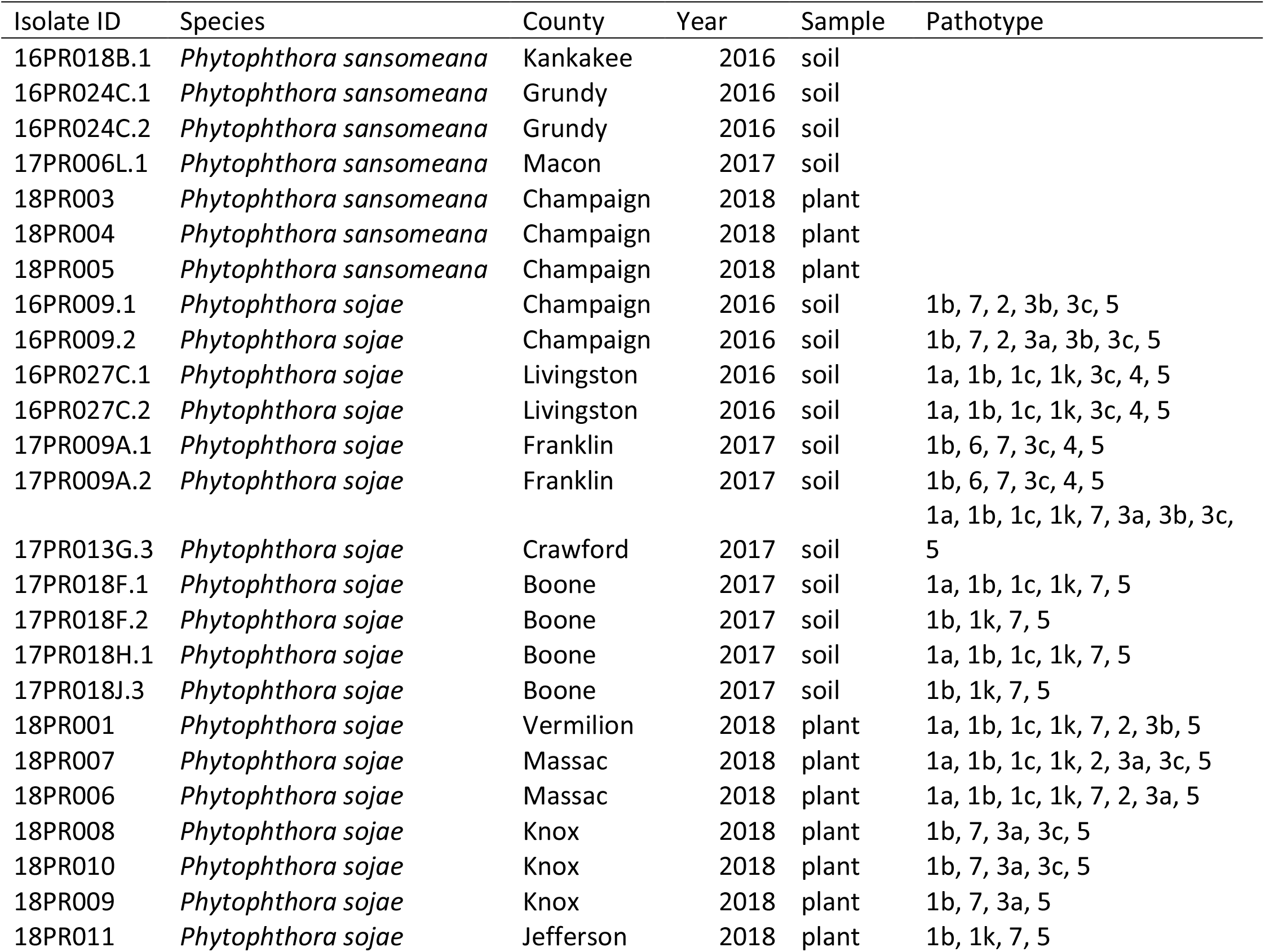
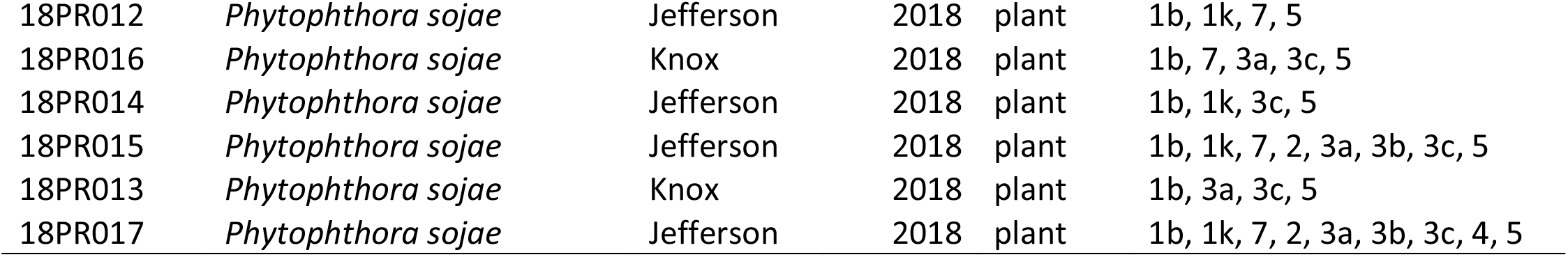
List of *Phytophthora* spp. isolates recovered from Illinois with the county and year collected, type of sample from where it was isolated, and pathotype identified in virulence assays.

Some soil samples contained both *Phytophthora* and *Pythium* spp. *P. sojae* was isolated with *Py. ultimum* var. *ultimum* and *Py. acanthophoron* from one of the soil samples. In another soil sample, it co-occurred with *Py. ultimum* var. *ultimum* and *Py. torulosum*. Neither *P. sojae* nor *Pythium* spp. were recovered from samples containing *P. sansomeana*.

### Virulence

Pathotypes for 24 *P. sojae* isolates were characterized by infecting 12 soybean differentials with known *Rps* genes and one susceptible control. Sixteen pathotypes were identified (Table 1). Only 8% of the isolates were virulent to *Rps*6 compared to 21% virulent to *Rps*4 and 25% to *Rps*3b (Figure 2A). All the isolates were virulent to *Rps*1b and *Rps*5. More than 50% of the isolates were virulent to *Rps*1k, and *Rps*7 and *Rps*3c. Thirty-three percent of the *P. sojae* isolates were virulent to *Rps*1a and *Rps*1c. The complexity of isolates ranged from four to nine, with a mean complexity of six. No isolate was virulent to all differential lines tested (Figure 2B).

**Figure 2.**
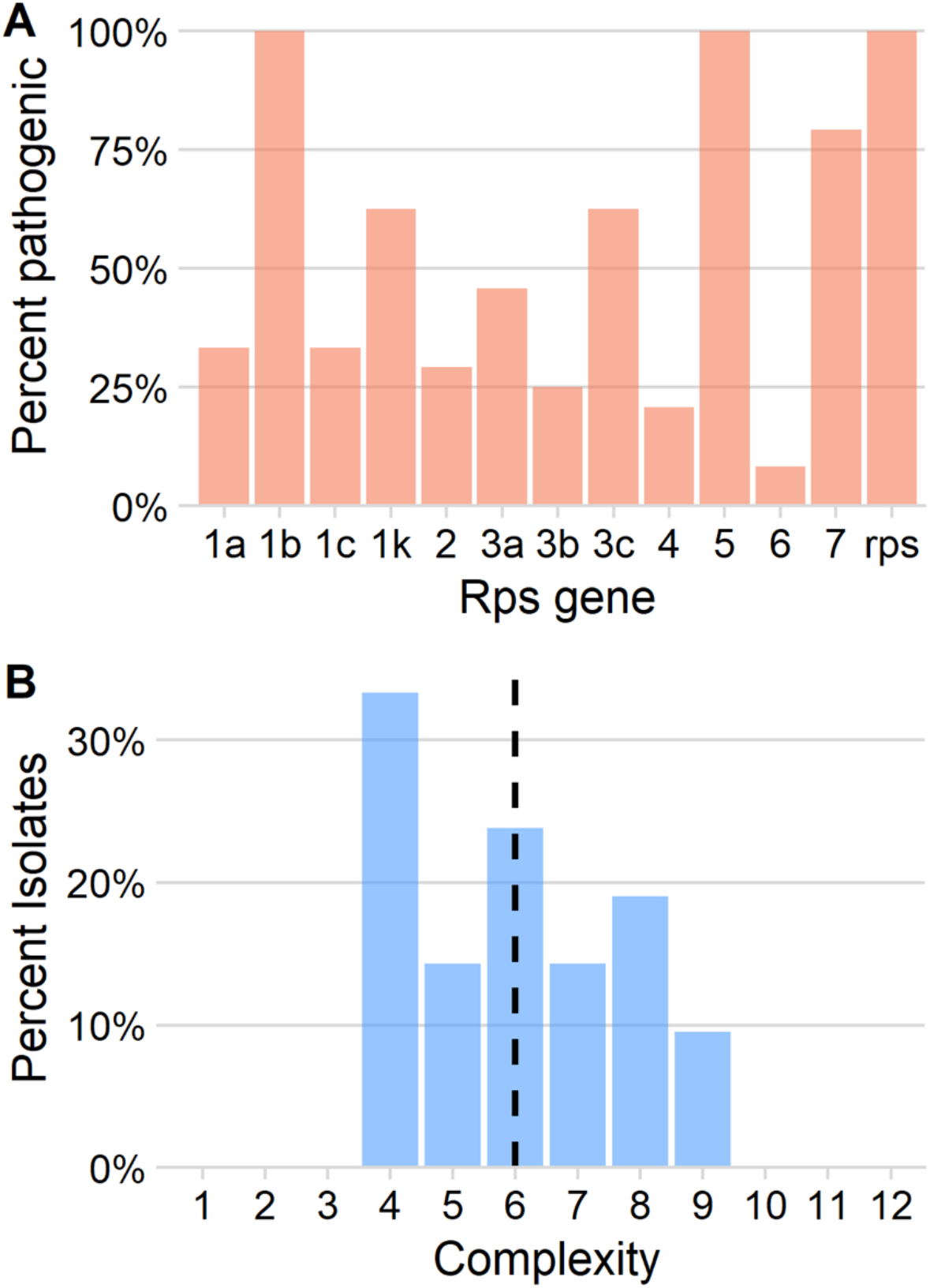
(A) Percentage of *P. sojae* isolates (n = 24) virulent to differentials with different *Rps* genes. (B) Frequency distribution of complexity (the number of *Rps* genes that an isolate is virulent to) of *P. sojae* isolates.

Of the 16 pathotypes of *P. sojae*, five were identified more than once and represented 80% of the total isolates (Table 1). Of these five pathotypes, only a single pathotype (1b, 1k, 7, 5) was recovered from more than one field. Within each field, pathotypes recovered ranged from one to four. Of the fields from which more than one *P. sojae* isolate was recovered, 43% contained isolates with a single pathotype, 43% with two pathotypes, and 14% with four pathotypes. In the fields where we recovered two pathotypes of *P. sojae*, isolate complexity ranged from four to eight, whereas complexity ranged from four to nine for the field with four pathotypes.

Most of the *P. sojae* differentials produced mixed results when inoculated with the *P. sansomeana* isolates, but some of the differentials performed consistently. None of the *P. sansomeana* isolates were able to cause disease on the *P. sojae* susceptible control rps (Williams). Interestingly, 43% of the *P. sansomeana* isolates were virulent to *Rps*3a (L83-570), 14% to *Rps*1c (Williams79), and 14% were virulent on both differentials.

### Aggressiveness of *Phytophthora* spp. on soybean

The aggressiveness of *Phytophthora* spp. isolates were assessed by infecting a susceptible cultivar and measuring root dry weight and shoot dry weight two weeks after germination. The average temperature of the greenhouse during the three experiments was 24.3°C (min: 20.89 - max: 29.6°C). Overall, *P. sojae* was more aggressive than *P. sansomeana* (Figure 3). Significant differences between species were observed for both root and shoot dry weight (*P* < 0.001). The root and shoot weight of *Phytophthora sojae* was significantly lower than the controls and *P. sansomeana*. No difference was observed between the controls and *P. sansomeana* for the root or shoot weight. On average, *P. sojae* reduced root weight by 54% and shoot weight by 48% compared to the control. This contrasts with *P. sansomeana*, which reduced root and shoot weight by 10% and 15%.

**Figure 3.**
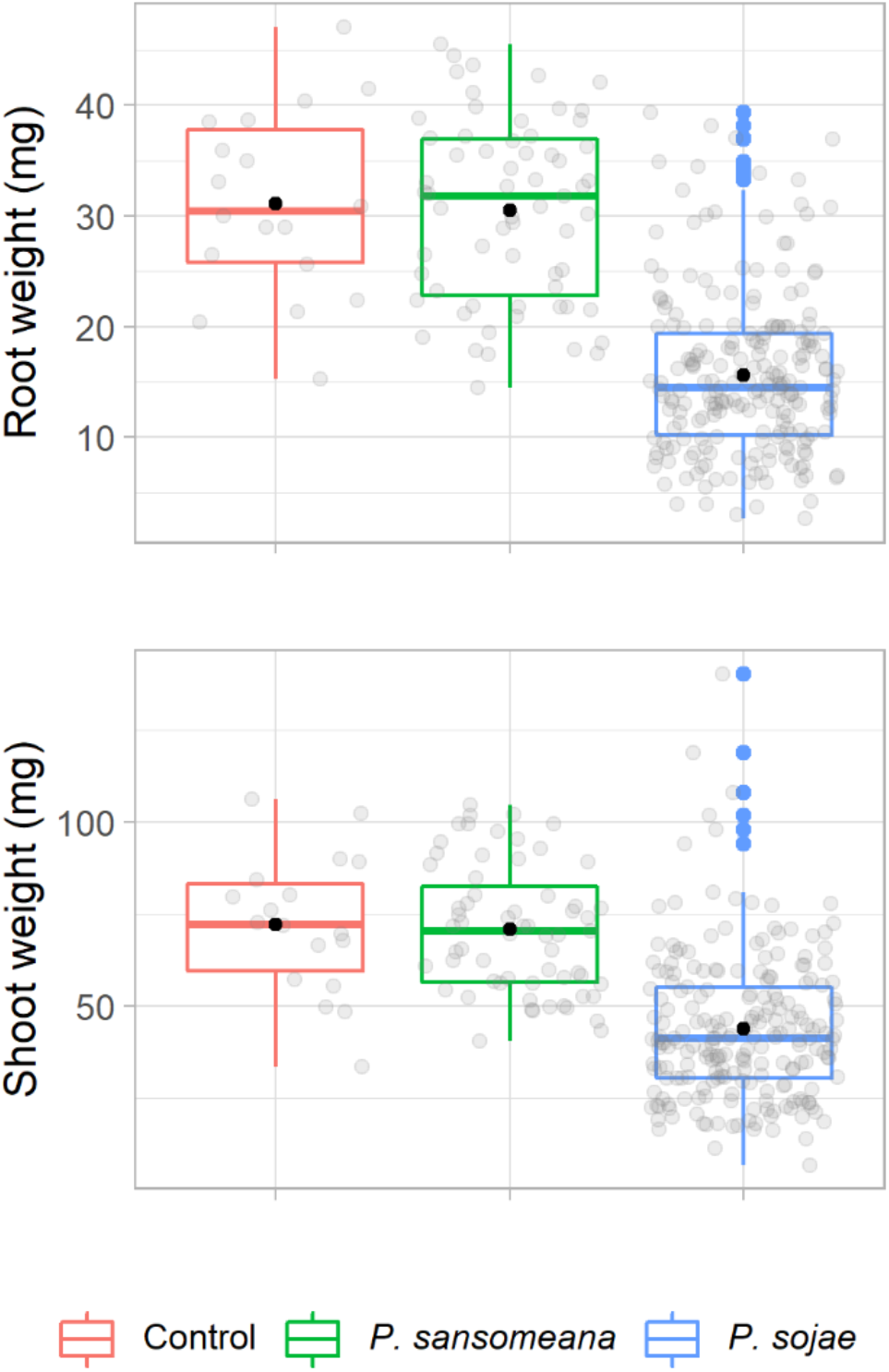
Distributions of root and shoot dry weights of the susceptible cultivar Sloan (no Rps), inoculated with isolates of *Phytophthora sansomeana* (n = 7) and *Phytophthora sojae* (n = 24). Center lines represent the medians, and solid black dots the means. Box limits indicate the 25th and 75th percentiles, and whiskers extend 1.5 times the interquartile range. Solid colored dots represent outliers.

Differences among isolates were observed for both root weight (*P* < 0.001) and shoot weight (*P* = 0.029). We observed a difference between the two controls for shoot weight, so isolate aggressiveness was compared against the mock control. For root weight, 23 *P. sojae* isolates (96%) were significantly different from the mock control, and only two were significantly less aggressive than the most aggressive isolate (Figure 4). The most aggressive isolate reduced root weight by 73%, while the least aggressive reduced root weight by 27%. Isolates not significantly different from the most aggressive isolate reduced root weight between 40 and 72%. The two isolates significantly different from the most aggressive isolate reduced root weight on average by 37%.

**Figure 4.**
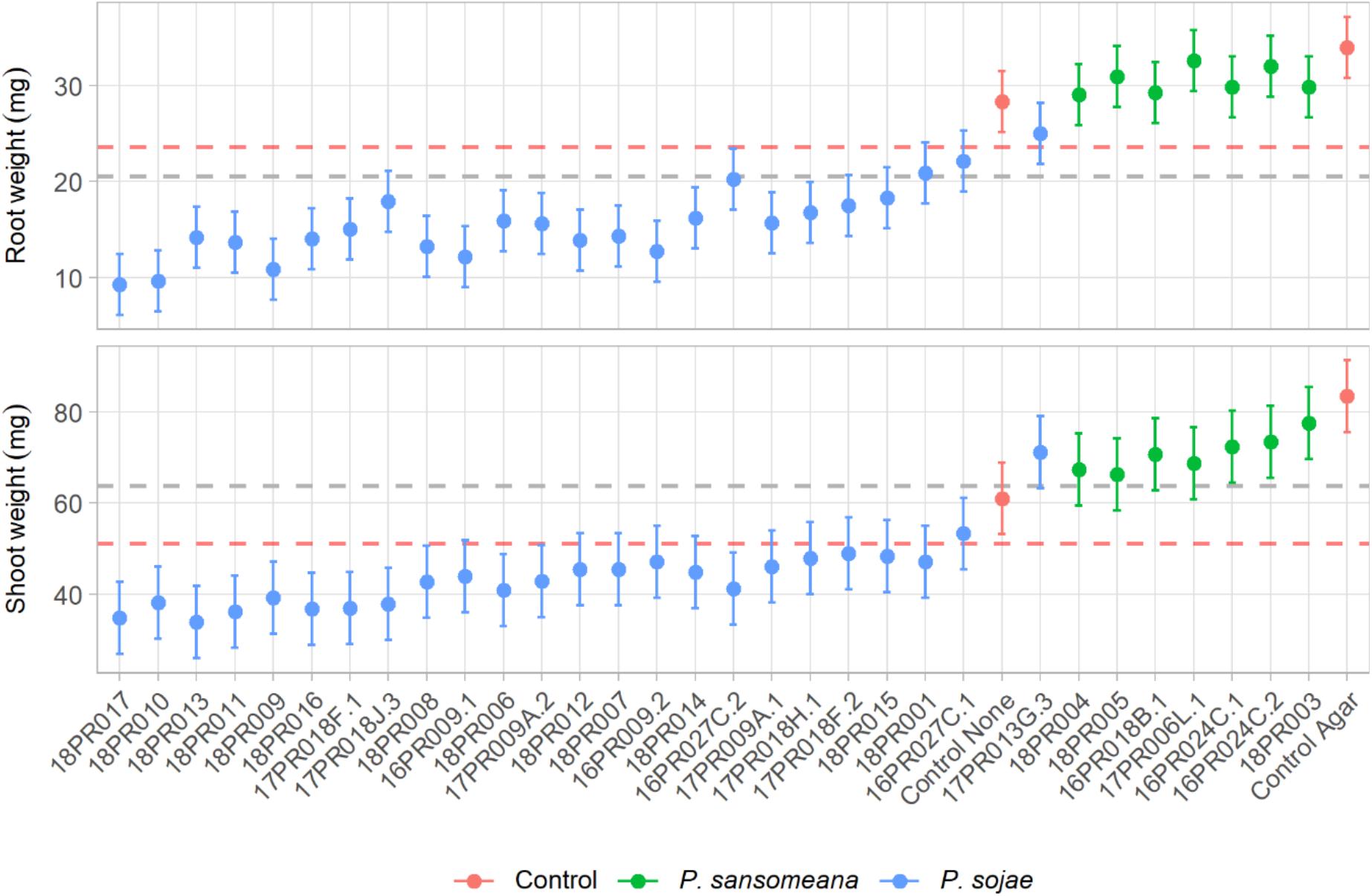
Mean root and shoot dry weight of the susceptible cultivar Sloan (*rps*) inoculated with isolates of *P. sansomeana* and *P. sojae*. Vertical lines represent the mean standard error. Means under the gray dotted line are significantly different from the mock control (agar) and means over the red dotted line are significantly different from the most aggressive isolate (18PR017) according to Tukey’s HSD test (*α* = 0.05).

For shoot weight, 22 *P. sojae* isolates (92%) were significantly different from the control but did not differ significantly from the most aggressive isolate (Figure 4). The most aggressive isolate reduced shoot weight by 59% compared to a 15% reduction by the least aggressive isolate. Isolates that did not differ significantly from the most aggressive isolate reduced shoot weight between 41% and 58%. The isolate for which significant differences were not observed compared with either the most aggressive isolate or the control reduced shoot weight by 39% on average.

None of the *P. sansomeana* isolates were significantly different from the mock control for root and shoot weight (Figure 4). All the *P. sansomeana* isolates differed significantly from the most aggressive *P. sojae* isolate for root and shoot weight. *P. sansomeana* isolates reduced root weight between 2% and 15% and shoot weight between 7% and 21%.

### Correlation between isolate complexity and aggressiveness of *P. sojae*

Pearson correlation analysis was performed to determine if the pathotype complexity of *P. sojae* isolates correlated with aggressiveness. Weak positive significant correlations were observed between isolate complexity with root weight (r = 0.18, *P* = 0.009) and shoot weight (r = 0.16, *P* = 0.015). Isolates with a higher complexity tended to be less aggressive (Figure 5). A moderate and significant correlation was detected between root and shoot weight (r = 0.54, *P* < 0.001).

**Figure 5.**
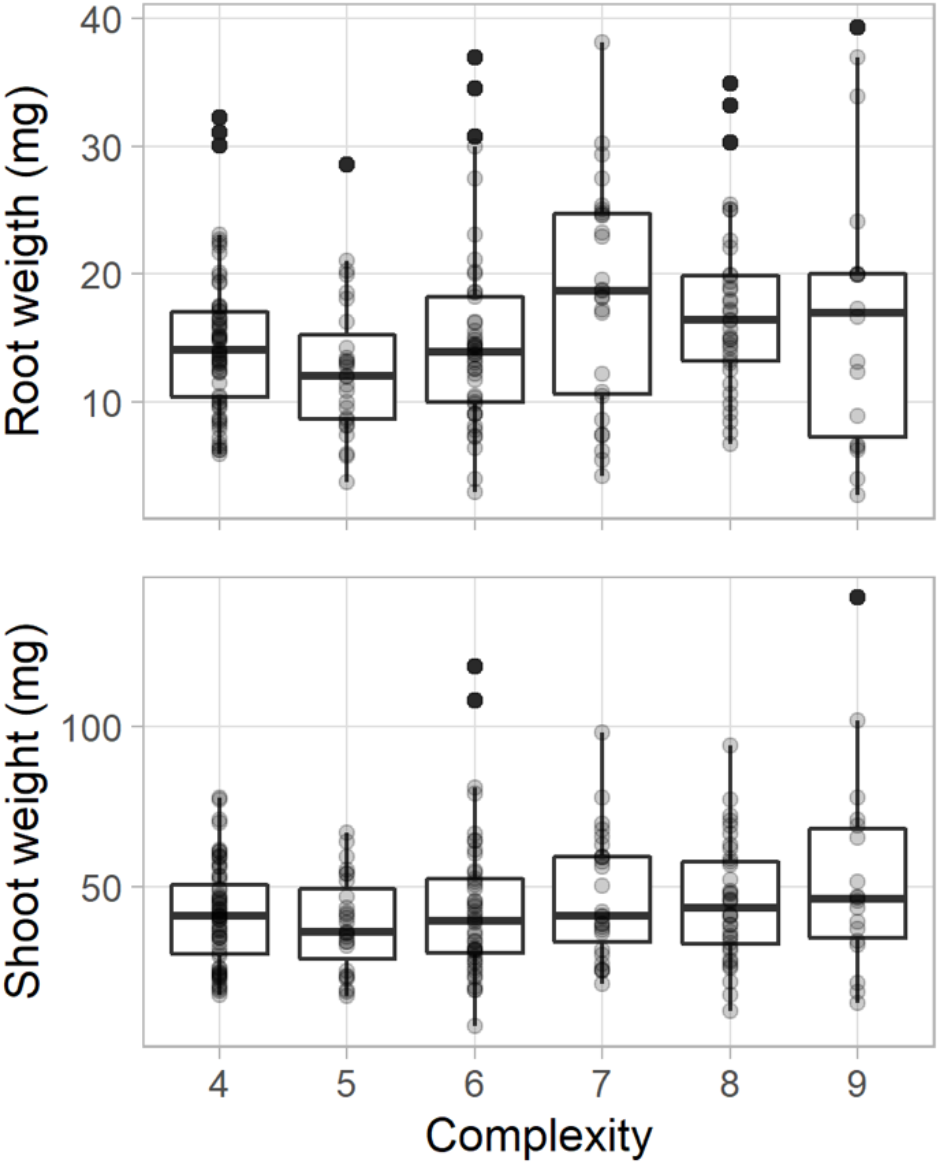
Distribution of root and shoot dry weight of the susceptible cultivar, Sloan (*rps*), inoculated with isolates of *P. sojae* (n = 24) grouped by complexity. Center lines represent the medians, and solid black dots the means. Box limits indicate the 25th and 75th percentiles, and whiskers extend 1.5 times the interquartile range. Pearson correlation between isolate complexity with root weight was r = 0.18 (*P* = 0.009) and isolate complexity with shoot weight was r = 0.16 (*P* = 0.015)

### Sensitivity of *Phytophthora* isolates to fungicides

All isolates of *P. sojae* (n = 9) and *P. sansomeana* (n = 7) were sensitive to the four fungicides tested (Figure 6), as indicated by a reduction in growth. On average, the highest EC_50_ values were observed for azoxystrobin (1.5 μg/ml), followed by mefenoxam (0.06 μg/ml), metalaxyl (0.05 μg/ml) and ethaboxam (0.02 μg/ml) indicating that ethaboxam was the most potent of the fungicides tested. Sensitivity to ethaboxam varied by species (Figure 6). On average, *P. sansomeana* was less sensitive to the tested fungicide active ingredients than *P. sojae* (0.023 μg/ml vs 0.008 μg/ml). *Phytophthora sojae* was more sensitive to azoxystrobin (1.2 μg/ml) than *P. sansomeana* (1.9 μg/ml). The response to mefenoxam and metalaxyl was similar for both species. Average EC_50_ values for mefenoxam were 0.060 μg/ml for *P. sojae* and 0.061 μg/ml for *P. sansomeana*. For metalaxyl, average EC_50_ values were 0.056 μg/ml for *P. sojae* and 0.051 μg/ml for *P. sansomeana*. Although no insensitive isolates were detected, the distribution of EC_50_ values shows that some isolates were less sensitive to all the fungicides tested (Figure 6). A *P. sansomeana* isolate (18PR005) had the highest EC_50_ values for all fungicides (Figure 6).

**Figure 6.**
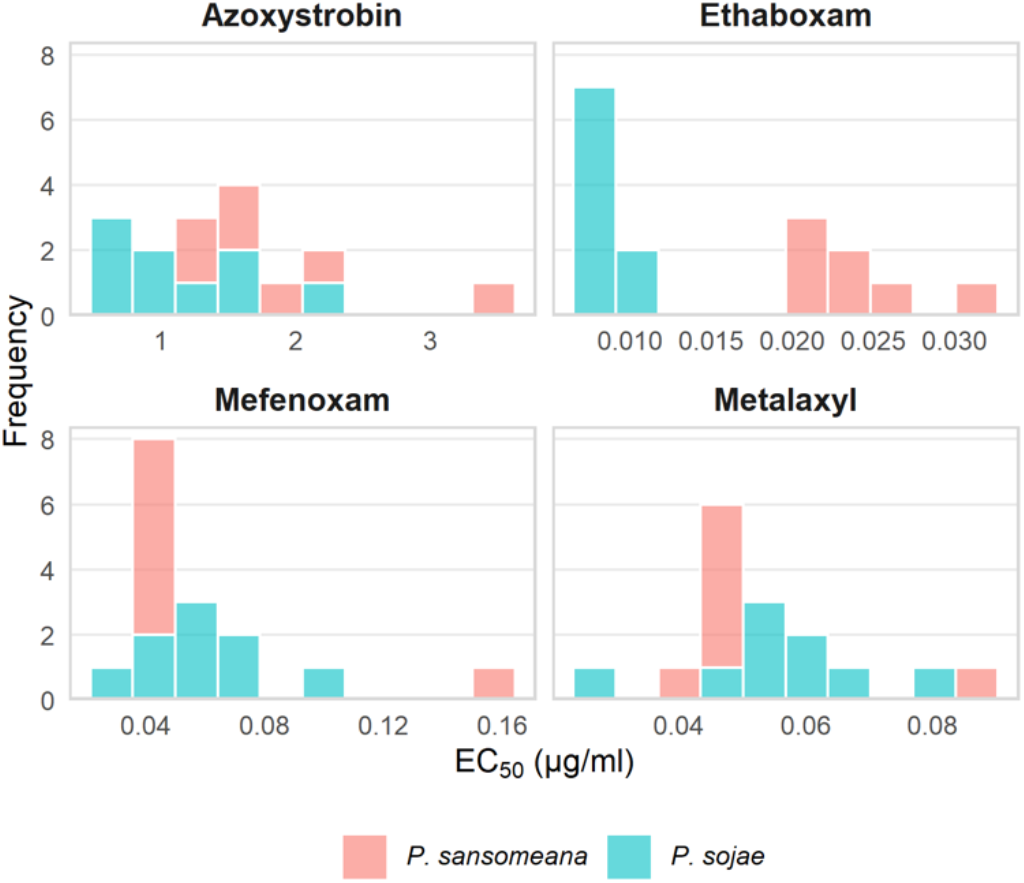
Distribution of EC_50_ values of *P. sojae* and *P. sansomeana* to technical grade azoxystrobin, ethaboxam, mefenoxam, and metalaxyl.

## Discussion

Monitoring the pathogen populations in a region is critical for informed disease management. Virulence characterization of *P. sojae* resulted in the identification of 16 pathotypes. In this study we present what is, to our knowledge, the first report of differences in aggressiveness between *P. sojae* isolates. In addition, we identified a possible reduction of aggressiveness on the more virulent isolates. Finally, we found no evidence in the *P. sojae* population for insensitivity to commonly used seed treatment active ingredients.

Isolates collected in this study were more complex than in past surveys suggesting an increase in virulence to several resistance genes. Compared to the 2012-2013 survey in Illinois (Dorrance et al. 2016), isolate mean complexity increased from 4.4 to 6.0. Put simply, the isolates in 2012-2013 were able to infect 4.4 differentials while the isolates in this study were able to infect 6 differentials on average. The most recent multistate collaboration survey reported a mean pathotype complexity of 6.6 for a subset of eleven isolates collected from Illinois soils between 2016 and 2017 (Hebb et al. 2022). Here we added *P. sojae* isolates collected from plant samples in 2018 and found an average mean complexity of six. Simple pathotypes, such as race 0 (virulent only to universal susceptible) and race 1 (*Rps*7), that were common in the past surveys (Leitz et al. 2000; Malvick and Gruden 2004; Dorrance et al. 2016), were not detected.

Based on the correlation between aggressiveness and isolate complexity, it could be inferred that incorporating more *Rps* genes into soybean should decrease the appearance of novel pathotypes of *P. sojae*. This assumes that aggressiveness of the isolate is a fitness-related trait. Despite the significant correlation between complexity and aggressiveness, the correlation was weak, and not all isolates with higher complexity were the least aggressive. If there was a cost for carrying additional virulence genes, the frequency of these genes should decrease over time in the pathogen population (Zhan and McDonald 2013). Future work should confirm the relationship between isolate aggressiveness and virulence to Rps genes and study the effect of isolate aggressiveness on quantitative disease resistance.

Reintroduction and rotation of the less commonly deployed genes, *Rps*3a and *Rps*6, has been proposed for long-term management of PRR (Dorrance et al. 2016; Yan and Nelson 2019; Matthiesen et al. 2021). Dorrance et al. (2016) reported that less than 10% of the isolates recovered in five of the 11 states surveyed for *P. sojae* pathotype diversity were virulent to these genes. However, in Illinois, more than 40% of isolates could overcome resistance from *Rps*3a. *Rps*6 was the most effective gene in the current study, with only two isolates (8%) virulent. Because these two isolates were recovered from the same field, *Rps*6 may still be effective in the majority of the fields in Illinois. Previous surveys in the state also reported only a few isolates that could overcome this gene (Leitz et al. 2000; Malvick and Gruden 2004). Cultivars with *Rps*6 are thus recommended to manage PRR; however, cultivars with *Rps*6 were uncommon in Illinois (Slaminko et al. 2010). Between 2003 and 2008, only two out of more than 3,000 cultivars that entered the University of Illinois Soybean Variety Testing Program contained *Rps*6 (Slaminko et al. 2010). Gene stacking of *Rps*3a and *Rps*6 with *Rps*1c or *Rps*1k has also been proposed (Dorrance et al. 2016). Gene stacking is also uncommon, but some companies have reported cultivars with *Rps*1c and *Rps*3a. Although more than 45% of our isolates could overcome *Rps*3a, only 13% of isolates recovered from two fields were virulent to *Rps*1c and *Rps*3a. Our results indicated a decrease in aggressiveness for the more complex pathotypes, suggesting that having stacks of resistance genes in soybean populations will reduce the damage by this pathogen. Deployment of two or more genes in soybean cultivars may be essential for the long-term management of PRR in Illinois.

*Phytophthora sansomeana* caused minor seedling damage under the same conditions that allowed significant damage from *P. sojae*. Also, it was recovered at lower frequencies compared to *P. sojae*. Optimum growth of *P. sansomeana* was reported to be between 25 and 27°C, and disease symptoms were observed at both 15 and 20°C in seedling assays (Hansen et al. 2009). This pathogen has been reported in multiple soybean-producing states, but few reports have studied the virulence and aggressiveness of this pathogen on soybean (Rojas et al. 2017a; Tande et al. 2020). Rojas et al. (2017a) reported this species as one of the most aggressive oomycetes in seed and seedling assays at 20°C, but disease did not manifest at 15°C. Navarro-Acevedo et al. (2021) inoculated cultivars with different resistance levels with four *P. sansomeana* isolates and reported different results depending on the isolate and cultivar. Only one isolate was virulent to Sloan, and the other two were virulent on cultivars Lorain (*Rps*1c) and Kottman (*Rps*1k, *Rps*3a) (Navarro-Acevedo et al. 2021). Both studies were performed in growth chambers compared to ours, which was conducted in the greenhouse. We used cultivar Sloan for aggressiveness evaluation, and average temperatures were close to 25°C with maximum temperatures up to 30°C. It has been reported that the aggressiveness of oomycetes varies significantly depending on the temperature (Matthienssen et al. 2016; Rojas et al. 2017a). This could further explain the absence of extensive disease in our results. As in our study, McCoy et al. (2021) reported inconsistent virulence results for *P. sansomeana* using the hypocotyl inoculation technique. Interestingly, for two of the differentials we obtained consistent results. Some of our isolates were able to defeat differentials with *Rps*1c and *Rps*3a. This suggests that there might be cross effect from some *Rps* genes towards *P. sansomeana*.

Both species were sensitive to all fungicides tested. Malvick and Grunden (2003) reported that *P. sojae* isolates recovered from Illinois were sensitive to mefenoxam and metalaxyl at concentrations <1 μg/ml and concluded that the risk of developing resistance in the state was low. Since then, no study in Illinois has been done reporting the sensitivity of *P. sojae* to these fungicides. Almost two decades later, our study does not show any resistance of *P. sojae* towards metalaxyl or mefenoxam. *P. sansomeana* was also sensitive to these fungicides, and no difference in sensitivity was observed compared to *P. sojae*. Our results agree with other studies that *P*. sansomeana and *P. sojae* are sensitive to fungicides (McCoy et al. 2021; Noel et al. 2019; Rojas et al. 2019). Ethaboxam is a newer compound, and other studies have found that both species are highly sensitive (Noel et al. 2019; Rojas et al. 2019). Ethaboxam was the most effective fungicide to reduce the growth of both species, but sensitivity varied by species. In this study, EC_50_ values for *P. sansomeana* (0.023 μg/ml) doubled the sensitivity of *P. sojae* (0.008 μg/ml). Noel et al. (2019) reported that *P. sansomeana* isolates recovered from soybean had EC_50_ values between 0.05 and 0.06 μg/ml, and one *P. sojae* isolate had an EC_50_ value of 0.02 μg/ml. Of the fungicides tested, azoxystrobin was the least effective at reducing fungal growth. *P. sansomeana* was less sensitive (1.9 μg/ml) to azoxystrobin than *P. sojae* (1.5 μg/ml). Shifts in sensitivity of *P. sansomeana* should be monitored since some oomycetes have an inherent resistance to ethaboxam, and there is a high risk of resistance selection to azoxystrobin (FRAC 2020; Noel et al. 2019).

In this study, we found that the virulence of *P. sojae* continues to increase in Illinois. Based on the virulence of the recovered isolates, we recommend the use of *Rps*6 and stacking of *Rps*6 and *Rps*3a with other resistance genes. Stacking is also supported by the reduction of aggressiveness with more virulence. *P. sojae* and *P. sansomeana* are still sensitive to the most commonly used active ingredients in seed treatments. As shown in other studies (e.g. Cerritos-Garcia et al. 2021) combining varieties with Rps genes, partial resistance, and seed treatments provide the most effective management for PRR.

## Acknowledgments

We thank Juan Pablo Granda, Adriana Noboa, Leigh Ann Fall, and Elizabeth Phillippi for technical support. Funding for this project provided by the North Central Soybean Research Program and the Department of Crop Sciences at the University of Illinois at Urbana-Champaign.

